# Phage growth rate is significantly affected by host-temperature and phage-temperature interactions but not by host-phage interactions

**DOI:** 10.1101/2025.05.06.651716

**Authors:** Paige Garrison, Daniel Padfield, Alec Martin, John Denby, Caroline Mincey, Jennifer L. Knies

## Abstract

Virus growth depends on host performance in any given environment and has a profound impact spanning fields from climate change to human health. We investigated the intrinsic growth rate of two natural G4-like bacteriophages on four clones of their host (*E*.*coli* strain C) that differed in their thermal performance, with significant differences in their maximum growth rate and optimum temperature. The intrinsic population growth rate of the G4-like phage isolates was measured on the four host clones at three temperatures, reflective of a low, optimum and high temperature for one of the phage. The phage’s intrinsic growth rate was significantly affected by host-temperature and phage-temperature interactions, but not by the interaction between host and phage. Across temperatures, phage growth rate was consistently highest on the host clone with the highest maximum growth rate, but was not, contrary to our expectations, lowest on the host with the lowest growth rate. At any given temperature, phage growth rate followed a directional trend with host growth rate, but this trend was not as apparent when looking between temperatures. Our work demonstrates a significant impact of temperature and the host on phage growth rate, but much work remains to be done to disentangle how variation in host performance impacts virus performance across temperatures.

## Introduction

Viruses, the most abundant organisms on earth (Hendrix *et al*., 1999; Mushegian, 2020), play a crucial role in global nutrient cycling (Suttle, 2005; Suttle, 2007) and infectious disease transmission, making the investigation of environmental effects on host-virus interactions essential for human health and global ecosystems (French & Holmes, 2020; Myers & Patz, 2009). Understanding how viruses regulate ecosystems as top-down parasites relies on understanding their adaptation to the environment and their host, with temperature being an important environmental factor (Harvell *et al*., 2002; Zimmerman *et al*., 2020). Although viruses generally exhibit greater resistance to thermal stress than their hosts (Madigan *et al*., 2018; Mojica & Broussaard, 2014), the ability of viruses to proliferate successfully within the host is influenced by temperature-dependent changes in host–virus interactions, partly driven by the host’s molecular response to temperature (Hochachka & Somero, 2002). By investigating these intricate dynamics, we can shed light on viral adaptation mechanisms that have implications for ecosystem functioning and disease transmission.

The temperature-dependence of virus survival and population growth is well established for diverse viruses, from early experiments on bacteriophages (Grozman & Suzuki, 1962), to recent work on viral pathogens including SARS-coV (Chan *et al*., 2011), malaria (Paaijmans *et al*., 2010), and influenza (Dalton *et al*., 2006). Naturally occurring lytic bacteriophages infecting the same host bacteria exhibit genetic variation in cardinal temperatures for growth (optimum, minimum, and maximum temperatures) (Knies *et al*., 2009; Rokyta *et al*., 2006). Additionally, Padfield *et al*. (2020) demonstrated cardinal temperatures shift for the host bacteria (*Pseudomonas fluorescens*) in the presence versus absence of its lytic bacteriophage, SBWΦ2. These studies and others have increased our understanding for the temperature-dependence of viral growth and host-virus interactions, but are limited by only considering how a single host affects - or is affected by - viral growth.

The interactions between viruses, their hosts, and temperature are difficult to predict due to the non-linear response of biological rates, such as growth rate, at the individual, population, and community level to changes in temperature (Huey & Kingsolver, 1989). Moreover, intraspecific (e.g. variation among host species physiology) or interspecific (e.g. virus infects a different host species) host variation may result in changes to the temperature response of host-virus interactions, as observed in viruses with diverse ectothermic hosts (Bryner & Rigling, 2011; Cheng *et al*., 2017; Shikano & Cory, 2015; Zouache *et al*., 2014). These interactions, termed genotype x genotype x environment (GxGxE), have been well documented in several pathogen-host systems, most prominently in microbial pathogens of agricultural plant pests (reviewed in Thomas & Blanford, 2003) but also in viruses such as the emerging Chikungunya virus (Zouache *et al*., 2014). Here we ask whether intraspecific variation in thermal performance of the bacteria Escherichia coli strain C affects the growth rate of two closely related phages across a wide temperature range. We predict phage growth will be highest on host strains that have the highest growth rates irrespective of thermal optimum of the host, but will increase within a single host as temperatures approach the optimum.

We confirm intraspecific variation in thermal performance for the host by measuring the growth of previously evolved *E. coli* clones across a wide temperature range. We then infected the host clones with G4-like phages known to be respectively more ‘warm’ and ‘cold’ adapted (Knies *et al*., 2009) at three temperatures. This allowed us to investigate how the growth of the phages depends on temperature and the host clone’s thermal performance. Future studies with more replicates and broader temperature ranges will be necessary to refine these findings and resolve more subtle host-phage dynamics.

## Materials and Methods

### Evolution of Bacteria Clones

To generate host (bacteria) clones with variation in their thermal performance, we used the standard laboratory host strain for the bacteriophage G4, *Escherichia coli* strain C (hereafter clone ANC). In prior experiments, the ancestral strain of the host was stored at -80°C, cultured at 37°C, and was used for phage experiments in batch culture (Holder and Bull, 2001; Knies *et al*., 2006; Knies *et al*., 2009). This ancestor was used to found a single bacterial population at each of three temperatures respectively representing a low, optimal, and high temperature for the bacteria: 20, 37 (the ancestral temperature), and 40°C. These populations evolved for 40 transfers in 10 mL Luria Broth (LB: 5 g yeast extract, 10 g Bacto tryptone, 5 g salt/1 L) at pH 7.5 and were maintained in 50 mL Erlenmeyer flasks placed in shaking incubators set to 200 rpm.

Evolving populations were propagated by transfers of 0.1 mL of stationary phase culture into 9.9 mL of fresh medium. Stationary phase was defined as an optical density (OD) at 600 nm (OD_600_) of greater than 1.8 on a spectrophotometer, which corresponded to 1-2×10^9^ colony forming units (CFU)/mL and was verified by plating. While density at which stationary phase is reached may be temperature dependent, the objective of experimental evolution was to generate clones with variation in their thermal performance, so any difference here is not important for this outcome. This resulted in transfers every 2-3 days at 20ºC and daily for the 37 and 42°C populations.

Between each transfer the population must grow ∼100-fold to reach stationary phase density, requiring log_2_100 = 6.64 generations per transfer. Failure to undergo this level of growth would result in rapid extinction. At 20 and 42°C, initial transfers (the first three to five) often required a greater transfer (up to 0.5 mL) to prevent extinction. A sample of each population was frozen at - 80°C every 5 transfers in 15% glycerol. After 40 transfers, one clone (colony) was chosen from each evolved population to yield clones *E*.*coli C*_*20E-40-I*,_ *E*.*coli C*_*37E-40-I*_, and *E*.*coli C*_*42E-40-I*_, hereafter clone 20E, clone 37E, and clone 42E.

To allow measures of phage growth on all four host clones simultaneously, as well as bacteria growth rate measures for all four clones at once, clones were cultured to stationary phase (2mL LB, shaking) at each experimental temperature and frozen in multiple single-use aliquots (100µL) in 15% glycerol at -80°C. Time to stationary phase was 24-48 hours depending on the temperature (48 hours at 20°C). Stationary phase was confirmed by optical density and plating as above. At temperatures above 45°C, clones did not achieve stationary phase so were not frozen.

### Phage culture conditions, intrinsic growth rate assays, and analysis

The G4 phage is a single-stranded DNA phage ideal for investigating effects of temperature on bacteria-phage interactions because its lifecycle and genome are well understood (Godson et al., 1978) and natural isolates of G4-like phages demonstrate variation in their optimum temperature and temperature range (Knies *et al*., 2006; Knies *et al*., 2009). The two G4-like phages (ID11 and ID12) used here were isolated from Idaho (Rokyta *et al*. 2006). Their thermal reactions norms on the ancestral host were described in Knies *et al*. (2009) and their *T*_*opt*_ were respectively 29.8ºC and 32.7ºC. As before, phage were cultured in LB supplemented with 2 mmol CaCl_2_ and on LB plates (15% agar). LB top agar (0.7% agar) was also supplemented with 2 mmol CaCl_2_. Phage were grown to high density on their standard laboratory *E*.*coli* strain C at room temperature (∼ 22ºC) (all phage formed well-defined plaques), then frozen in single use aliquots at -20°C in LB broth supplemented with 2 mM CaCl_2_ and 40% glycerol.

Phage growth rate assays were performed at three temperatures (20, 28, and 40°C) on all four bacterial clones (clone ANC, clone 20E, clone 37E, and clone 42E). These three temperatures represented low to high temperatures for these phages which respectively had minimum, optimum, and maximum temperatures of ∼22, 30 and 37°C (ID11) and ∼ 24, 33, and 41°C (ID12) (Knies *et al*. 2009). For this experiment, we modified the Knies *et al*. (2009) phage phage growth rate assays to be performed in a small(er) volume (100µL), and by performing the experiments in a thermocycler, allowed for temperature control, and for growth rate to be measured on all four host clones at once. Four biological replicate assays were performed at each temperature, but some were dropped for the analysis as a growth rate could not be calculated as the phage had gone extinct (below our detection level) likely due to a long lysis time at the extreme temperatures at 20 and 40°C. All assays used the previously frozen hosts, with single-use aliquots of each clone thawed, diluted to a lower density, then distributed in 90 µL aliquots into 0.2 mL strip tubes and cultured at the experimental temperature in a thermocycler until exponential phase (see Methods below, mean CFU/90 µL = 7.67x 10^6^).

Bacteria were only thawed once. Phage stocks were adjusted to 10^5^ phage/mL and 10µL, ∼ 10^3^ phages, were added to this exponentially growing culture. The low ratio of phage to bacteria (< 0.002) allowed the observed growth rate to be the maximum possible. After 45 minutes the culture of phage and bacteria was treated with 5 µL chloroform to lyse the bacterial cells. Phage numbers at the start (N_0_) and end (N_45_) of the assay were determined by plating on the standard *E*.*coli* strain C host, incubating the plates at room temperature for 16-24 hours and counting the number of plaque forming units (PFUs/mL). Intrinsic rate of population growth (*r*) per hour was calculated as *r* = [ln(N_45_/N_0_)]/0.75. Intrinsic growth rate for the phage is referred to as phage growth rate hereafter.

We used an analysis of variance (ANOVA) to assess the effects of phage genotype, host clone, temperature, and all possible interactions on the phage growth rate. Host clone, phage genotype, and temperature were treated as categorical variables. The *anovan* function in Matlab (version R2020b) was used. The residuals for the fitted model were normally distributed (Anderson-Darling test statistic = 0.27, h =0). A multiple comparison test (Matlab function *multcompare*) was used to compare phage growth rate averaged across temperatures between bacteria clones.

To test the relationship between phage growth rate and host growth rate, we did Pearson’s correlation on the phage growth rate and the predicted host growth rate of each clone from the best fit of the thermal performance curve at 20, 28, and 40ºC

### Measuring bacteria growth across temperatures

Bacteria clone growth rates were measured under conditions mimicking the phage growth rate assays (100μL volume without shaking) and were not meant to mimic the adaptive environment for the bacteria populations. Variation in thermal performance between host clones was confirmed by measuring growth curves for each clone across their full thermal tolerance to estimate the optimum and maximum temperature of growth. To ensure the exponential growth phase was observed, pilot experiments between 20 and 48°C were performed to determine the experiment length needed to capture exponential growth and the temperature range for the experiment.

Growth curves for the bacteria clones (clones ANC, 20E, 37E, and 42E) were measured using the frozen stocks (see above) at each of 10 temperatures: 20, 25, 28, 32, 37, 42, 45, 48, 52, and 58°C. The temperatures were chosen to include several temperatures above the optimum and to span a wide temperature range (Pawar *et al*., 2016) for an accurate estimate of the thermal reaction norms. For temperatures above 45ºC, the 45°C frozen stocks were used. Similar to Garcia *et al*. (2018) and Padfield *et al*. (2020), growth was measured in 96 well plates.

Experiments were performed in flat-bottomed 96 well plates (Corning, Costar) covered with sealing film (Thermo electron, Mylar plate sealers) to prevent evaporation. The bacterial clones were thawed from a -80°C stock (described above) and each aliquot used in a single growth rate assay. The cells were returned to low density by diluting the cells 1:10 to ∼1-2 × 10^7^ cells per 100 μL. Seven to ten samples (100 μL each) per clone were put into individual wells (one row per clone) and a corresponding 10 wells at the top and bottom row of the plate served as controls, each with 100 μL of LB media. We saw no contamination in our control wells. The first and last column of each plate were left empty. The plate was placed in a temperature-controlled incubator and allowed to incubate at the experimental temperature for half an hour before the first measurement to avoid measurements affected by air bubbles. Optical density (OD_600_) was measured as a proxy for bacterial density using a plate reader. Readings of OD_600_ were taken with the sealing film off every 20 minutes for up to five hours, except at 42 and 45°C which was 3 hours 40 minutes because of faster growth at these temperatures. Time out of the incubator was minimized and cultures were plated at the beginning of each experiment to verify cell density and the absence of contamination.

### Calculating exponential growth rate for bacteria

We performed a sliding window analysis in R using lm()to fit a linear model to the log_10_(OD_600_) as the response variable and time as a predictor over all sets of three consecutive time points (∼1 hour) for each technical replicate (a single well) for each clone at each temperature. Exponential growth rate (hereafter referred to as bacterial growth rate), was the maximum slope from these fits (total of 9-12 fits) (Madigan *et al*., 2018; Padfield *et al*., 2020). For the measurements at 32°C, we restricted the growth rate estimate to the first 200 minutes of growth as OD_600_ measurements became biologically unrealistic. We considered a modelling approach to estimating growth rate using the R package ‘nlsMicrobio’ (Baty & Delignette-Muller, 2013), but could not capture the observed increase in growth rate with temperature.

Individual estimates of growth rate from each technical replicate were then passed to the thermal reaction norm model fitting process.

### Fitting temperature performance curves (TPCs) to bacterial growth rate

Many models have been used to describe the continuous asymmetric shape of TPCs (reviewed in Kontopoulos *et al*. (2023), Low-Décarie *et al*. (2017), and Padfield *et al*. (2021)), but most studies still only fit single models and do not justify their choice. We considered equations (8, 9, 10) described in Low-Décarie *et al*. (2017) that all allowed direct estimation of our key ecologically relevant parameters of interest: *T*_*opt*_. However, preliminary analyses with our data showed equation 8 was unable to capture the curve asymmetry and equation 9 struggled to model the shape of the curve around the optimum. The best fitting model was equation 10, as assessed by the visual fits of the models to the data and the AIC score for each model to each clone (Supplemental Table 2). We therefore used equation 10 in Low-Décarie *et al*. (2017) (based on Eppley (1972), see also Norberg (2004) and Thomas *et al*. (2012)).

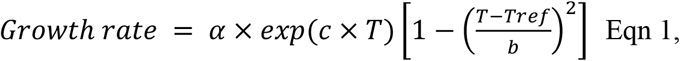

where the growth rate of the bacteria clones increases with temperature according to α × *exp*(*c* ×*T*), which is an envelope function (Eppley, 1972) that puts an upper limit on maximum growth rate. The decline of growth rate is controlled by the right-hand side of Eqn 1: 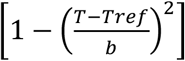. The overall shape of the TPC depends on thermal niche (*b*) and the rise in growth rate as given by *a* and *c*. To estimate *a, b, c*, and *T*_*ref*_ for each bacteria clone, model fits were performed using all the individual technical replicates of growth rate for each clone in Matlab (version R2020b) using the *lsqcurvefit* function, with *a* and *c* constrained to have a minimum value of zero, *T*_*ref*_ constrained to between zero and 100 and, *b* constrained to between ten and 100.

We used the fitted parameters from Eqn 1 to calculate the cardinal growth parameters for each clone: the optimum temperature (*T*_*opt*_), maximum growth rate (*R*_*max*_), and upper critical temperature (*T*_*max*_). We did not estimate the lower critical temperature because accurate estimates of the lower critical temperature require very low growth rates (Low-Décarie *et al*., 2017) and these were not practical to measure. The optimum temperature was solved by numerical optimization using the *fminbnd* function in Matlab. The maximum growth rate was the growth rate at the optimum temperature. The upper critical temperature was estimated from the roots of Eqn 1, where the growth rates switch from positive to negative.

### Evaluating the difference between clones’ cardinal growth parameters

We estimated whether the cardinal growth parameters differed between clones using bootstrapping. To account for uncertainty in our estimates of the cardinal growth parameters we used a parametric bootstrapping approach as in Listman *et al*. (2016) to bootstrap the residuals (Wehrens *et al*., 2000) of the fitted model (Eqn 1). We performed 1000 residual bootstraps, a procedure in which the residuals of the best model fit are randomly ‘reassigned’ to the predicted values (each of which corresponds to a growth rate measurement), thereby generating a slightly different thermal reaction norm. For each iteration, we refitted Eqn 1 to estimate the model parameters (*a, c, T*_*ref*_, and *b*) and calculated the cardinal growth parameters: *T*_*opt*_, *T*_*max*_, and *R*_*max*_. From these bootstraps, we calculated 95% confidence intervals for each parameter. If the 95% confidence intervals of the trait did not overlap between two clones (Schenker & Gentleman, 2001), we concluded that the trait differed significantly between the two clones.

## Results

### Evolved *E. coli* clones differ in their optimum temperatures and maximum growth rates

The TPCs for all host bacteria clones showed the typical unimodal shape (Huey & Kingsolver, 1989) (Figure 1a and Figure S2). Crucially, as measured under conditions mimicking the phage growth rate assays, clones had significant differences in their thermal performance (Figure 1). Consistent with evolution at low temperature, clone 20E had a lower *T*_*opt*_ of 38.7 °C (95% confidence intervals [CI]: 36.4–40.7 °C) compared to 42.21°C (95% CI: 40.60-43.67°C) for clone ANC. Moreover, clone 37E evolved a higher *T*_*opt*_ of 45.20°C (95% CI: 43.87-45.99°C) than the 20E clone and the ANC clone (Figure 1b). Consistent with evolution at a stressful (high) temperature, clone 42E had the lowest *R*_*max*_ of 0.23 hr^-1^ (95% CI: 0.22-0.24 hr^-1^), but we saw no evolution in its *T*_*opt*_ (Figure 1). The ancestral clone (ANC) had the highest *R*_*max*_, with all clones being significantly different except clones 20E and 37E. *T*_*max*_ ranged from 57.61 to 59.09 °C and was not significantly different between any pairs of clones (Figure 1d).

**Figure 1.**
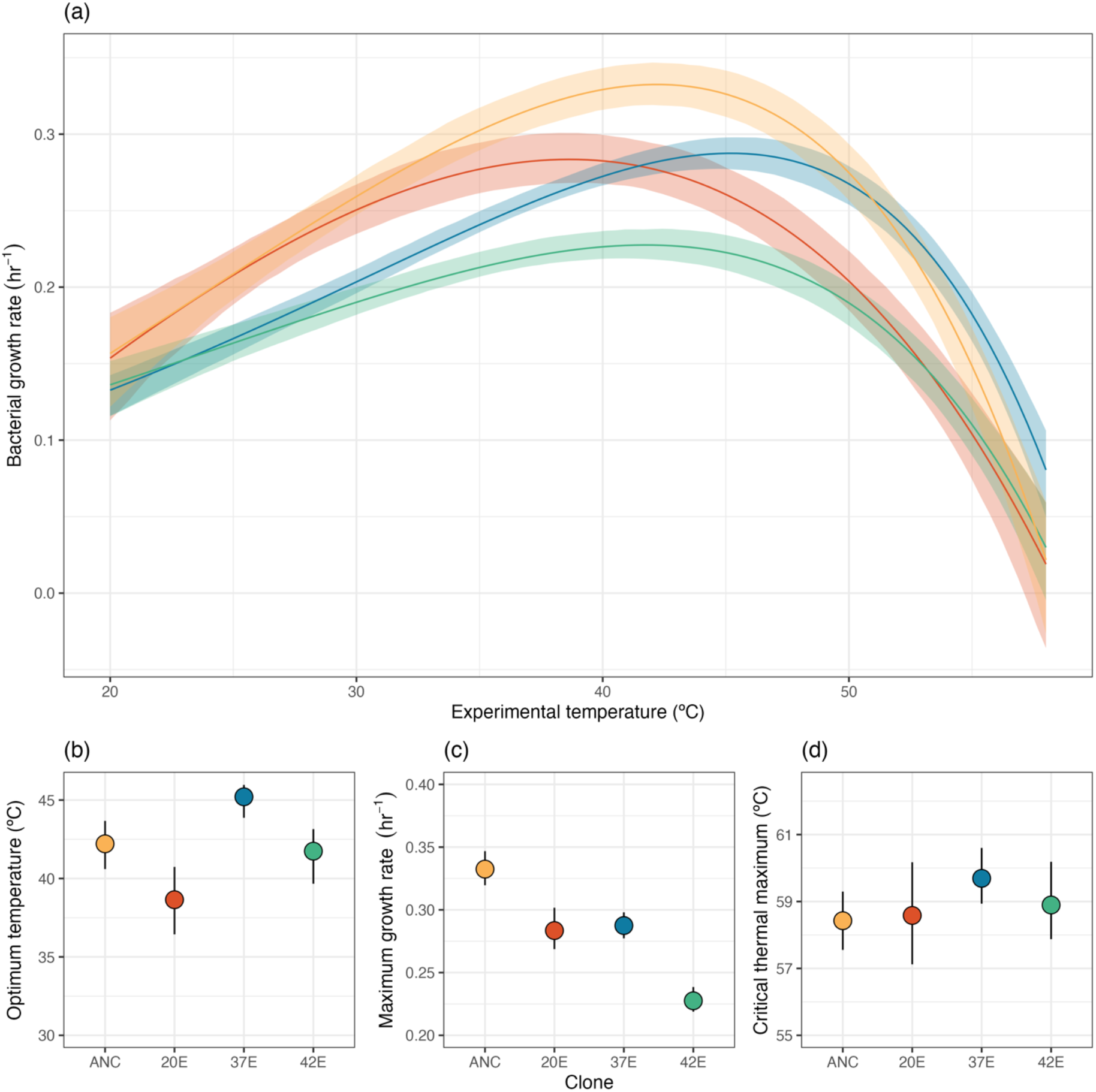
Thermal performance of *E. coli* clones after experimental evolution at different temperatures. (a) *E. coli* clone growth showed a unimodal response to temperature, but differences were seen from the ancestor (orange) after experimental evolution at 20 (red), 37 (blue) and 42ºC (green). Experimental evolution at different temperatures resulted in some differences in (b) optimum temperature and (c) maximum growth rate, but no differences in (d) critical thermal maximum. Non-overlapping confidence intervals indicate significant differences. In (a) the solid line represents the prediction from the best model fit for each clone and shaded bands represent 95% confidence intervals of predictions. In (b-d) points and lines represent the prediction from the best model fit and 95% confidence intervals of the estimated parameters.

### Phage growth differs by phage identity, *E. coli* clone and temperature

Growth rate of both phages were highly temperature dependent, ranging from an intrinsic rate of population growth of over 10 (representing a >22,000 increase in phage count) at 28ºC to negative growth at 20ºC (representing a decrease in phage counts from the beginning of the assay). Unfortunately, this meant that some replicates (12 of the 96) had phage titres below (or in one case, above) our detection limit. Removing the points reduced the power of our analyses.

Phage growth rate increased concurrent with an increase in temperature from 20 to 28°C across all host clones, but when moving from 28 to 40ºC, growth rate decreased for the cold-adapted phage (ID11) (Figure 2a), but increased or remained unchanged for the hot-adapted phage (ID12) (Figure 2b) (temperature x phage interaction: F_(2,60)_=44.06, p<0.0001). Host clone significantly impacted phage growth rate (F_(3,60)_=10.56, p < 0.0001), with growth rate being highest on the ANC clone, intermediate on the 42E and 20E clones, and worst on the 37E clone (Supplemental Table 1). The effect of host clone on phage growth rate varied by temperature (temperature x host interaction: F_(6,60)=_3.68, p= 0.0036) . The phage-host and phage-host-temperature interactions were not significant (Table 1).

**Figure 2.**
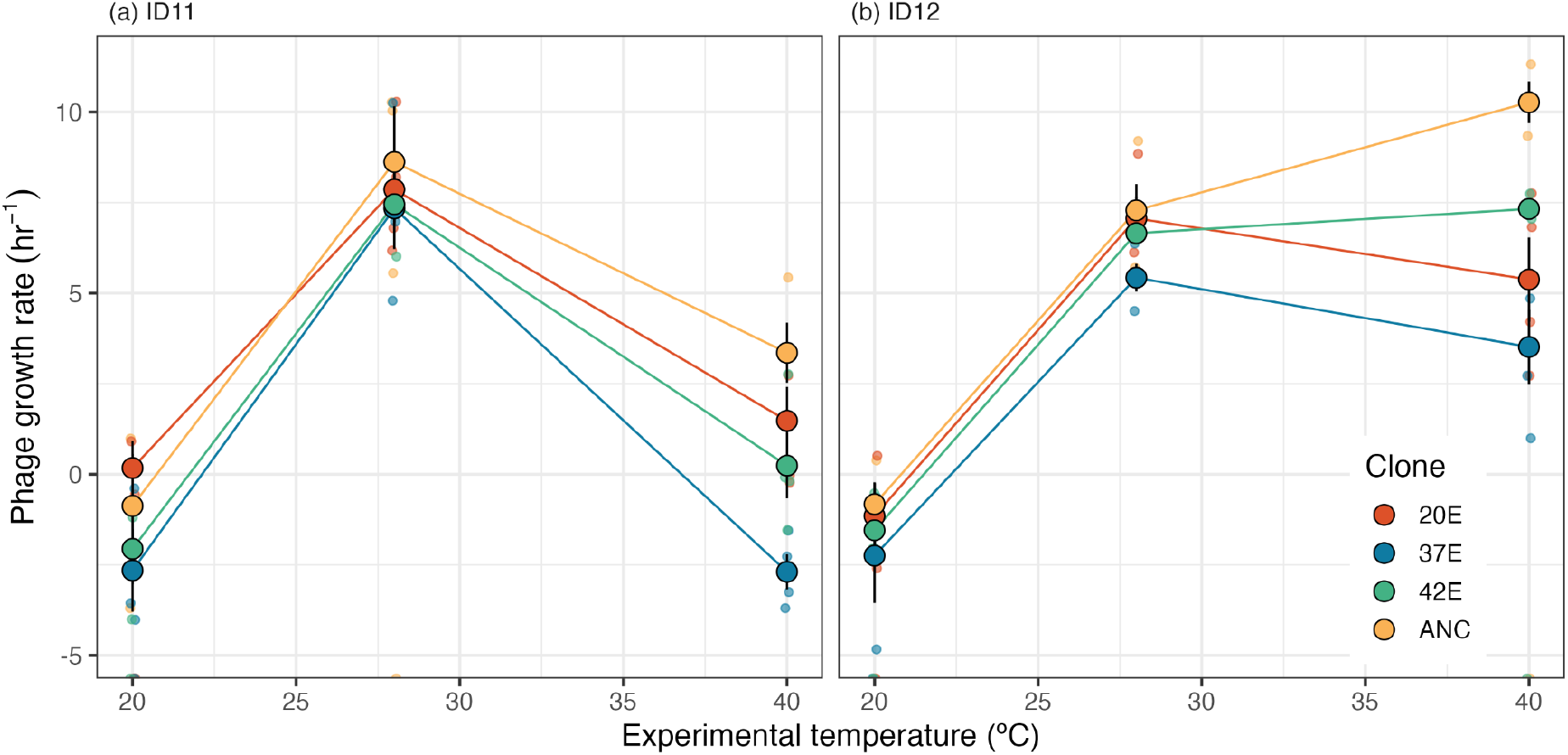
Phage growth rates across temperature and between host bacteria clones. Growth rate was measured on the four different *E. coli* clones across three temperatures for (a) the cold-adapted phage ID11 and (b) the warm-adapted phage ID12. Large points represent means of technical replicates and error bars represent standard errors. Small points are the individual technical replicates.

**Table 1.** ANOVA describing the effect of phage genotype, host clone, and temperature on phage growth rate. Significant predictors are indicated in bold.

| Source of Variation | SS | d.f. | MS | F | p-value |
| --- | --- | --- | --- | --- | --- |
| Host | 108.39 | 3 | 36.13 | 13.26 | **<0.0001** |
| Phage | 50.52 | 1 | 50.52 | 18.54 | **0.0001** |
| Temp | 965.79 | 2 | 482.89 | 177.25 | **<0.0001** |
| Host*temp | 60.09 | 6 | 10.015 | 3.68 | **0.0036** |
| Phage*temp | 211.49 | 2 | 110.74 | 40.65 | **<0.0001** |
| Phage*host | 7.67 | 3 | 2.56 | 0.94 | 0.43 |
| Phage*host*temp | 9.30 | 6 | 1.55 | 0.57 | 0.75 |
| Error | 163.46 | 60 | 2.72 |  |  |
| Total | 1592.69 | 83 |  |  |  |

To test whether phages have higher growth rates when bacteria are growing faster, a Pearson correlation was performed for each phage between the phage growth rate at each temperature and predicted host growth rate at that temperature. Across all temperatures, there is a strong positive correlation between predicted bacterial growth rate and phage growth rate for ID12 (Pearson correlation coefficient = 0.80, P-value = 0.0019), but it is much weaker for the more cold-adapted ID11 (0.25, P-value = 0.43), due to the decline in growth rate at 40ºC. A Pearson correlation was also performed between phage and bacterial growth for each phage genotype within each temperature individually. A positive relationship exists between phage and bacterial growth for both phage genotypes, with Pearson correlations ranging from 0.44 to 0.92 for individual phage-temperature combinations (Figure 3).

**Figure 3.**
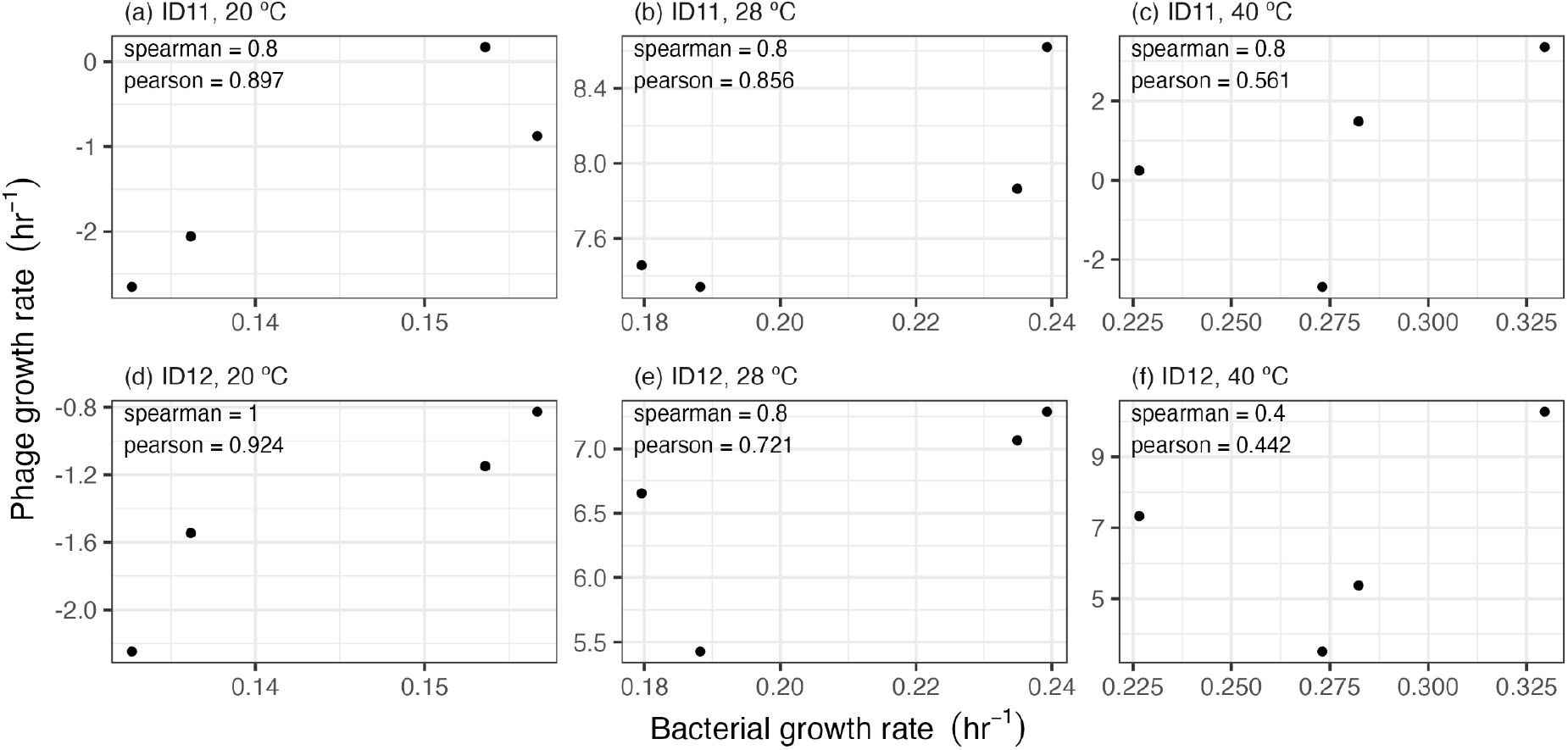
Phage growth rates as a function of predicted host bacterial growth rate at each experimental temperature. Phage data are mean growth rates for two to four biological replicates per temperature. Host growth rate is predicted growth rate for each clone from the fitted Eqn 1 parameters at 20, 28, and 40°C. Each panel represents a unique phage-temperature combination and shows growth rate for that phage on each host clone at the listed temperature.

## Discussion

How intraspecific variation in host thermal performance impacts viral thermal performance remains an unanswered question. In bacteria-phage research, previous work on the effect of temperature has been done on single host clones, but variation in host thermal performance may, in turn, alter phage thermal performance. Research in this area is limited and our study is admittedly small but it begins to point out ways to perform these experiments, shows the importance of host clone will probably be easy to detect, and suggests that detecting interactions may be more difficult and will need careful experimental consideration of temperatures for host and phage including prospective power analyses. We generated clones of a standard bacterial host (*E. coli* strain *C*) with intra-specific variation in TPCs, with clones having significant differences in their *T*_*opt*_ and *R*_*max*_ (Figure 1). Phage growth rate depended on an interaction between the phage and temperature and on an interaction between host clone and temperature (Figure 2 and Table 1). Consistent with our predictions, we found phage growth was highest on host strains that have the highest growth rates irrespective of thermal optimum and increased within a single host population as temperatures approached the optimum (Figure 3). The relationship between phage growth rate and predicted host growth rate was stronger for the more hot-adapted phage (ID12) than the more cold-adapted phage (ID11).

There were several interesting results that are secondary to the main question of how intraspecific variation in thermal performance of hosts impacts the growth rate of parasites across temperatures. During our experiments, we surprisingly found that the ancestral clone (ANC) had the highest *R*_*max*_. While interesting, this likely reflects differences in selection imposed during short-term assays measuring growth in 96 well plates, and those imposed during batch culture. For example, while the ancestral E. coli is predicted to have a *T*_*opt*_ of 42.21°C from our TPCs, ancestral *E. coli* struggled initially at 42ºC in batch culture, which was reflected in the TPC of the the 42E clone C with a lower *R*_*max*_.

This new dataset is the first work to look at intraspecific variation of both the bacteria and the phage. It is valuable and provides future directions for research, but our inferences are limited by our experiments, which are always a trade-off between what would be ideal and what is feasible. Firstly, we are limited by the variation in thermal performance of both the phage and bacterial host. We used experimental evolution to produce variation in host thermal performance, with evolution temperatures having a wide range of 20ºC. Despite this, differences in thermal performance of our evolved clones differed by only 6.5ºC for *T*_*opt*,_ which is less than seen for interspecific variation (Alster *et al*., 2017; Barberán *et al*., 2017; Bronikowski *et al*., 2001).

Although we still observe significant differences in phage growth across clones, we would expect to see larger effects if the difference between bacteria host response was larger. Using three replicates for each phage growth rate assay is commonly used (Knies *et al*., 2006; Knies *et al*., 2009) but we were limited by some of our four replicates having phage counts below our level of detection. This is because the phage are unable to infect the host at these temperatures, or because phage are still undergoing replication during the 45 minutes of the assay. The removal of this data may have limited our statistical power to detect any significant three-way interaction between phage, host, and temperature. Measuring phage growth across more temperatures and measuring more replicates within each temperature will increase our ability to understand the importance of intraspecific variation on host-parasite interactions. Finally, our TPCs analyses use estimates of growth rate from microplate readers as the response variable to fit curves to. We do not propagate uncertainty across analysis steps, but the average error for individual growth rate estimates were small and did not systematically change across temperatures. For these reasons, this lack of error propagation is unlikely to change our results, but we look forward to method development that makes it easier to propagate uncertainty across an analytical pipeline.

We propose two areas of focus for future work. First, investigate whether phage-host and phage-host-temperature interactions affect phage growth rate by increasing the replication, and associated statistical power, for phage growth rate assays. Additionally, focus the experiments on temperatures at the upper limit for the phage where differences between phage genotypes are largest, making interactions easier to detect. Second, assess whether intraspecific host variation in TPCs affects the phage’s cardinal temperature for growth by measuring phage growth rate at higher resolution across a wide temperature range. Phage growth rate depends on key phage traits-principally attachment rate, burst size, and the latent period (Madigan *et al*., 2018). These traits are affected by temperature, among other external factors (Jończyk *et al*., 2011). Measuring these traits at different temperatures will provide a mechanism for how changes in host thermal performance affects phage growth rate. Phage burst size (or calculated growth rate) is established as being positively correlated with host growth rate (or dilution rate) and inversely correlated with latent period at single temperatures in other phage systems (Hadas *et al*., 1997; Golec *et al*., 2014; Mancuso *et al*., 2018; Nabergoj *et al*., 2018; Santos *et al*., 2014). We plan to extend our experiments to a larger set of naturally isolated G4-like phages, allowing us to explore the broader ecological implications of virus performance on different host clones.

## Conclusion

Overall, our work contributes to the growing body of knowledge on temperature-dependent host-viral interactions, which are sometimes excluded from studies on thermal reaction norms because of the unique parasitic life cycle of viruses (Malusare *et al*., 2022). By highlighting the complexities of these dynamics, we pave the way for more targeted research on virus adaptation mechanisms, ultimately enhancing our understanding of ecosystem dynamics and disease transmission in a changing global environment.

## References

Alster, C.J., Weller, Z. D., & von Fischer, J.C. (2017). A meta-analysis of temperature sensitivity as a microbial trait. Global Change Biology, 24(9), 4211–4224. 10.1111/gcb.14342

Barberán, A., Velazquez, H.C., Jones, S., & Fierer, N. (2017). Hiding in plain site: Mining bacteria species records for phenotypic trait information. mSphere, 2(4), e00237–17. 10.1128/mSphere.00237-17

Baty, F. & Delignette-Muller, M. L. (2013). nlsMicrobio: data sets and nonlinear regression models dedicated to predictive microbiology.

Bronikowsk, A., Bennett, A.F., & Lenski, R.E. 2001. Evolutionary adaptation to temperature. VIII. Effects of temperature on growth rate in natural isolates of Eschericia coli and Salmonella enterica from different thermal environments. Evolution, 55 (1), 33–40. 10.1111/j.0014-3820.2001.tb01270.x

Bryner, S.F. & Rigling, D. (2011). Temperature-dependent genotype-by-genotype interaction between a pathogenic fungus and its hyperparasitic virus. The American Naturalist, 177(1), 65–74. 10.1086/657620

Chan, K.H., Malik Peiris, J.S., Lam, S.Y., Poon, L.L.M., Yuen, K.Y., & Seto, W.H. (2011). The effects of temperature and relative humidity on the viability of the SARS Coronavirus. Advances in Virology, 2011, 734690. 10.1155/2011/734690

Cheng, K., Van de Waal, D.B., Niu, X.Y., & Zhao, Y.J. (2017). Combined effects of elevated pCO2 and warming facilitate cyanophage infections. Frontiers in Microbiology, 8, 1096. 10.3389/fmicb.2017.01096

Dalton, R.M., Mullin, A.E., Amorim, M.J., Medcalf, E., Tiley, L.S., & Digard, P. (2006). Temperature sensitive influenza A virus genome replication results from low thermal stability of polymerase-cRNA complexes. Journal of Virology, 3, 58. 10.1186/1743-422X-3-58

Eppley, R. W. Temperature and phytoplankton growth in the sea. (1972). Fishery Bulletin, 70 (4), 1063–1085.

French, R.K. & Holmes, E.C. (2020). An Ecosystems Perspective on Virus Evolution and Emergence. Trends in Microbiology, 28(3), 165–175. 10.1016/j.tim.2019.10.010

García, F.C., Bestion, E., Warfield, R., & Yvon-Durocher, G. (2018). Changes in temperature alter the relationship between biodiversity and ecosystem functioning. Proceedings of the National Academy of Sciences,115(43), 10989–10994. 10.1073/pnas.1805518115

Godson, G.N., Barrell, B.G., Staden, R., & Fiddes, J.C. (1978). Nucleotide sequence of bacteriophage G4 DNA. Nature, 276(5685), 236–247. 10.1038/276236a0

Golec, P., Karczewska-Golec, J., Łoś, M., & Węgrzyn, G. 2014. Bacteriophage T4 can produce progeny virions in extremely slowly growing Escherichia coli host: comparison of a mathematical model with the experimental data, FEMS Microbiology, 351(2), 156–161. 10.1111/1574-6968.12372

Groman, N.B. & Suzuki, G. (1962). Temperature and lambda phage reproduction. Journal of Bacteriology, 84(3), 431–437. 10.1128/jb.84.3.431-437.1962

Hadas, H. Einav, M., Fishov, I., & Zaritsky, A. 1997. Bacteriophage T4 development depends on the physiology of its host Escherichia coli. Microbiology, 143, 179–185. 10.1099/00221287-143-1-179

Harvell, C.D., Mitchell, C.E., Ward, J.R., Altizer, S., Dobson, A.P., Ostfeld, R.S., & Samuel, M.D.(2002). Climate warming and disease risks for terrestrial and marine biota. Science, 296, 2158–2162. 10.1126/science.1063699

Hendrix, R.W., Smith, M.C., Burns, R.N., Ford, M.E., & Hatfull, G.F. (1999). Evolutionary relationships among diverse bacteriophages and prophages: all the world’s a phage. Proceedings of the National Academy of Science1s, 96(5), 2192–2197. https://doi.org/10.1073/pnas.96.5.2192

Hochachka, P. & Somero, G. (2002). Biochemical adaptation: mechanism and process in physiological evolution. Oxford University Press.

Huey, R. B. & Kingsolver, J. G. (1989). Evolution of thermal sensitivity of ectotherm performance. Trends in Ecology and Evolution, 4 (5), 131–135. 10.1016/0169-5347(89)90211-5

Jończyk, E., Kłak, M., Międzybrodzki, R. & Górski, A. (2011) The influence of external factors on bacteriophages—review. Folia Microbiologica, 56, 191–200. 10.1007/s12223-011-0039-8

Kontopoulos, D.G., Sentis, A., Daufresne, M., Glazman, N., Dell, A.I., & Pawar, S. (2024). No universal mathematical model for thermal performance curves across traits and taxonomic groups. Nature Communications, 15, 8855. 10.1038/s41467-024-53046-2

Knies, J. L., Izem, R., Supler, K.L., Kingsolver, J.G., & Burch, C.L. (2006). The genetic basis of thermal reaction norm evolution in lab and natural phage populations. PloS Biology, 4(7), e201. 10.1371/journal.pbio.0040201

Knies, J.L., Kingsolver, J.G., & Burch, C.L. (2009). Hotter is better and broader: thermal sensitivity in a population of bacteriophages. The American Naturalist, 173(4), 419–430. 10.1086/597224

Listmann, L., LeRoch, M., Schlüter, L., Thomas, M.K., & Reusch, T.B. (2016). Swift thermal reaction norm evolution in a key marine phytoplankton species. Evolutionary Applications, 9(9), 1156–1164. 10.1111/eva.12362

Low-Décarie, E., Boatman, T.G., Bennett, N., Passfield, W., Gavalás-Olea, A., Siegel, P., & Geider, R.J. (2017). Predictions of response to temperature are contingent on model choice and data quality. Ecology and Evolution, 7(23), 10467–10481. 10.1002/ece3.3576

Madigan, M.T., Bender, K.S., Buckley, D.H., Sattley, W.M, & Stahl, D.A. (2018). Brock Biology of Microorganisms (15th edition). Pearson.

Malusare, S., Zilio, G. & Fronhofer, E.A. (2022). Evolution of thermal performance curves: A meta-analysis of selection experiments. Journal of Evolutionary Biology, 36(1), 15–28. 10.1111/jeb.14087

Mancuso, F., Shi., J., & Malik D.J. 2018. High throughput manufacturing of bacteriophages using continuous stirred tank bioreactors connected in series to ensure optimum host bacteria physiology for phage production. Viruses, 10(10), 537. 10.3390/v1120100537

Mojica, K.D.A. & Broussard, C.P.D. (2014). Factors affecting virus dynamics and microbial host-virus interactions in marine environments. FEMS Microbiology Ecology, 89(3), 495–515. 10.1111/1574-6941.12343

Mushegian, A.R. (2020). Are There 1031 Virus Particles on Earth, or More, or Fewer? Journal of Bacteriology, 202(9), e00052–20. 10.1128/JB.00052-20

Myers, S.S. & Patz, J.A. (2009). Emerging Threats to Human Health from Global Environmental Change. Annual Review of Environment and Resources, 34 (1), 223–252. 10.1146/annurev.environ.033108.102650

Nabergoj, D., Modic, P., & Podgornik, A. (2018). Effect of bacterial growth rate on bacteriophage population growth rate. MicrobiologyOpen, 7(2), e558. 10.1002/mbo3.558

Norberg, J. (2004). Biodiversity and ecosystem functioning: A complex adaptive systems approach. Limnology and Oceanography, 49(4, part 2), 1269–1277. 10.4319/lo.2004.49.4_part_2.1269

Paaijmans, K.P., Blanford, S., Bell, A.S., Blanford, J.I., Read, A.F., & Thomas, M.B. (2010). Influence of climate on malaria transmission depends on daily temperature variation. Proceedings of the National Academy of Sciences, 107(34), 15135–15139. 10.1073/pnas.1006422107

Padfield, D., Castledine, M., & Buckling, A. (2020). Temperature-dependent changes to host– parasite interactions alter the thermal performance of a bacterial host. The ISME Journal, 14(2), 389–398. 10.1038/s41396-019-0526-5

Padfield, D., O’Sullivan, H., & Pawar, S. (2021). rTPC and nls.multstart: A new pipeline to fit thermal performance curves in R. Methods in Ecology and Evolution, 12(6), 1138–1143. 10.1111/2041-210X.13585

Pawar, S., Dell, A.I., Savage, V.M., & Knies, J.L. (2016). Real versus artificial variation in the thermal sensitivity of biological traits. The American Naturalist, 187(2), E41–E52. 10.1086/684590

Rokyta, D. R., Burch, C.L., Caudle, S.B., & Wichman, H.A. (2006). Horizontal gene transfer and the evolution of microviridcoliphage genomes. Journal of Bacteriology, 188(3), 1134–1142. 10.1128/JB.188.3.1134-1142.2006

Santos, S.B., Carvalho, C., Azeredo, J., & Ferreira, E.C. 2014. Population dynamics of a Salmonella lytic phage and its host: implications of the host bacterial growth rate in modelling. PLoS One, 9(7), e102507. Erratum in: PLoS One. 201510(8), e0136007. 10.1371/journal.pone.0102507.

Schenker, N. & Gentleman, J. (2001). On judging the significance of differences by examining the overlap between confidence intervals. The American Statistician, 55(3), 182–186. 10.1198/000313001317097960

Shikano, I. & Cory, J.S. (2015). Impact of environmental variation on host performance differs with pathogen identity: Implications for host-pathogen interactions in a changing climate. Scientific Reports, 5, 15351. 10.1038/srep15351

Suttle, C.A. (2005). Viruses in the sea. Nature, 437(7057), 356–361. 10.1038/nature04160

Suttle, C.A. (2007). Marine viruses-major players in the global ecosystem. Nature Reviews Microbiology, 5(10), 801–812. 10.1038/nrmicro1750

Thomas, M.B. & Blanford, S. (2003). Thermal biology in insect–parasite interactions. Trends in Ecology and Evolution, 18(7), 344–350. 10.1016/S0169-5347(03)00069-7

Thomas, M.K., Kremer, C.T., Klausmeier, C.A., & Litchman, E. (2012). A global pattern of thermal adaptation in marine phytoplankton. Science, 338 (6110), 1085–1088. 10.1126/science.1224836

Wehrens, R., Putter, H., & Buydens, L.M.C. (2000). The bootstrap: a tutorial. Chemometrics and Intelligent Laboratory Systems, 54 (1), 35–52 . 10.1016/S0169-7439(00)00102-7

Zimmerman, A.E, Howard-Varona, C., Needham, D.M., John, S.G., Worden, A.Z, Sullivan, M.B., Waldbauer, J.R., & Coleman, M.L. (2020). Metabolic and biogeochemical consequences of viral infection in aquatic ecosystems. Nature Reviews Microbiology, 18(1), 21–34. 10.1016/S0169-5347(03)00069-7

Zouache, K., Fontaine, A., Vega-Rua, A., Mousson, L., Thiberge, J.M., Lourenco-De-Oliveira, R., Caro, V., Lambrechts, L., & Failloux, A.B. (2014). Three-way interactions between mosquito population, viral genotype and temperature underlying chikungunya virus transmission potential. Proceedings of the Royal Society B, 281(1792), 20141078. 10.1098/rspb.2014.1078

