## Supplemental Material for "Phage growth rate is significantly affected by host-temperature and phage-temperature interactions but not by host-phage interactions"

**Supplemental Figure 1**

A clone of the bacteria *E.coli* genotype strain *C* was used to found three populations – one at 20°C, one at 37°C, and one at 42°C. Each population was propagated through 40 serial transfers at its selective temperature. In each serial transfer cycle, the population was diluted 1:100 and went through $\log_2 100 = 6.64$ generations of binary fission per cycle. A single isolate was used from the 40th transfer of each population for subsequent experiments.

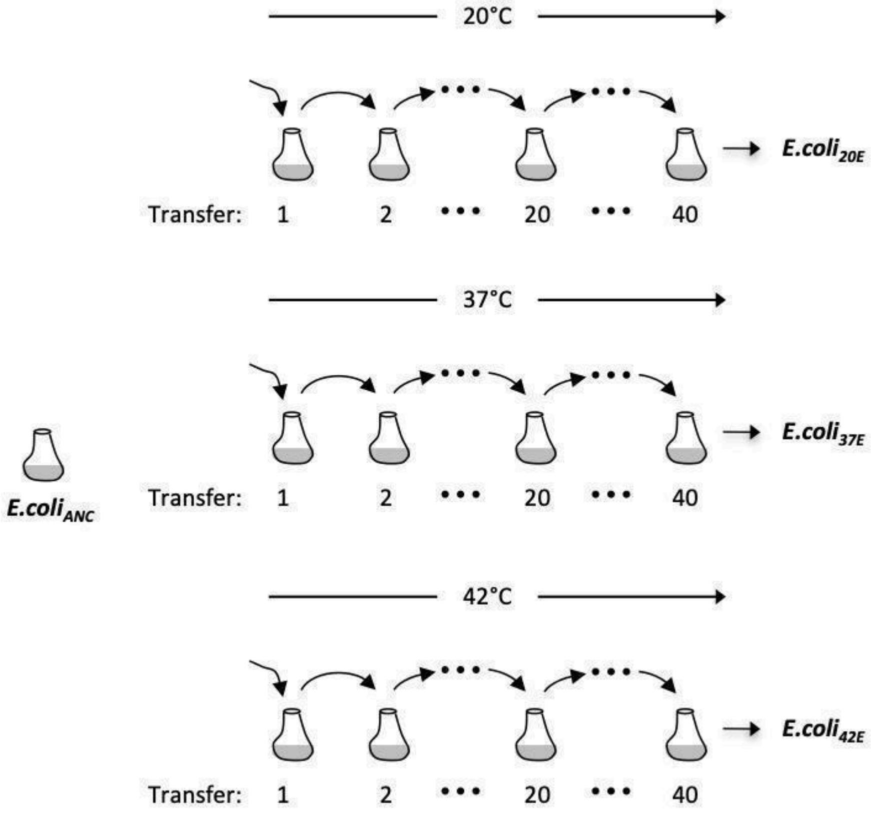


.

**Supplemental Figure 2. Thermal performance curves for repeated measures of bacterial growth rate across temperature.** Points represent repeated measures of exponential growth rate from one experiment at each temperature (See Methods for growth rate protocol). The data at at 32℃ were constrained (see Methods). Solid lines represent the mean prediction and shaded bands represent the 95% confidence interval of predictions.

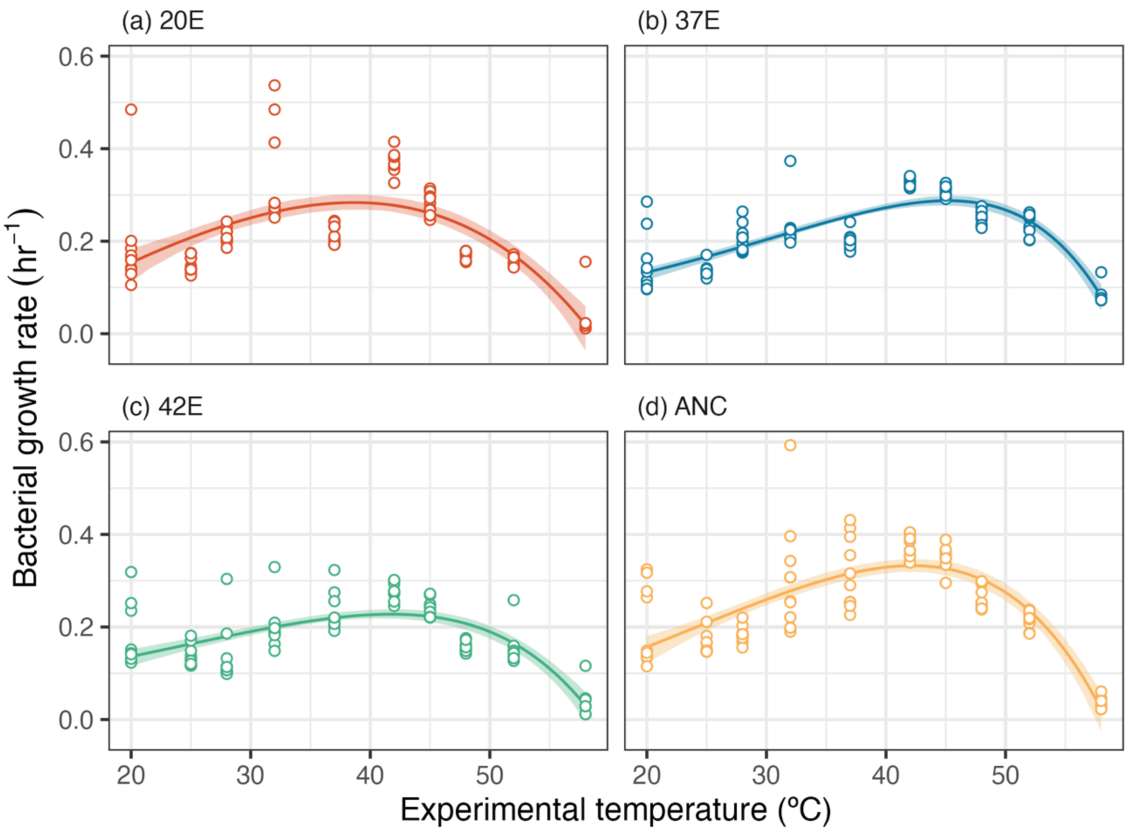

**Supplemental Table 1.** Phage intrinsic growth rate varies between pairs of bacteria clones. A multiple comparison test was performed after the analysis of variance because clone had a significant main effect on phage growth rate in the anova (see Table 1). Significance is indicated in bold.

| Clone one | Clone two | Lower Limit | Difference | Upper Limit | p-value |
|---|---|---|---|---|---|
| ANC | 20E | -0.22 | 1.17 | 2.56 | 0.13 |
| ANC | 37E | 1.83 | 3.19 | 4.55 | **<0.0001** |
| ANC | 42E | 0.25 | 1.65 | 3.00 | **0.014** |
| 20E | 37E | 0.66 | 2.02 | 3.38 | **0.0012** |
| 20E | 42E | -0.92 | 0.45 | 1.83 | 0.82 |
| 37E | 42E | -2.91 | -1.47 | -0.22 | **0.016** |

**Supplemental Table 2.** AIC scores for models considered for fitting thermal reaction norms to bacteria growth rate data. The number of parameters in each model is indicated by **k.** The lowest AIC score for each clone is indicated in **bold.**

| Model | Citation | k | Clone | logL | AIC |
|---|---|---|---|---|---|
| $Rate = a \cdot exp\left[-0.5 \cdot \left(\frac{[T - T_{ref}]}{b}\right)^2\right]$ | Eqn 8 in Low-DeCarie (2017) | 3 | ANC | 116.1005 | -226.2011 |
| | | | 20E | 110.96 | -215.91 |
| | | | 37E | 149.86 | -293.72 |
| | | | 42E | 139.45 | -272.91 |
| $Rate = a \cdot exp\left[-0.5 \cdot \left(\frac{abs[T - T_{ref}]}{b}\right)^c\right]$ | Eqn 9 in Low-DeCarie (2017) | 4 | ANC | 116.95 | -225.89 |
| | | | 20E | 111.45 | -214.90 |
| | | | 37E | 158.61 | -309.22 |
| | | | 42E | 144.87 | -281.73 |
| $Rate = a \cdot exp(c \cdot T)\left[1 - \left(\frac{[T - T_{ref}]}{b}\right)^2\right]$ | Eqn 10 in Low-DeCarie (2017) | 5 | ANC | 123.95 | **-245.91** |
| | | | 20E | 112.40 | **-222.79** |
| | | | 37E | 166.06 | **-330.13** |
| | | | 42E | 146.98 | **-291.96** |
